# Combining pupillometry and drift-diffusion models reveals auditory category learning dynamics

**DOI:** 10.1101/2024.04.16.589753

**Authors:** Jacie R. McHaney, Casey L. Roark, Matthew J. McGinley, Bharath Chandrasekaran

**Affiliations:** Northwestern University; University of New Hampshire; Baylor College of Medicine

## Abstract

Pupillometry and drift-diffusion models (DDM) have emerged as powerful tools to offer insights into learning processes and decisional dynamics. Specifically, pupillary dilation can serve as a metric of arousal and cognitive processing, and DDMs examine processes underlying perceptual decision-making using behavioral accuracies and response times. Methodological constraints have complicated the combination of the two methods in the study of learning. DDMs require precise response times, yet pupillary responses are slow and are impacted by motor movements. Here, we developed a learning task that separately optimized measurement of behavioral response times and pupil dilation during learning. We modified a standard learning paradigm to include trials with and without timing delays between stimulus presentation and response. Delayed trials optimized measurement of the pupil, while immediate trials optimized behavioral response times. We tested whether decision-making processes estimated with DDMs could be recovered from delayed trials where pupil response measurement was optimized. Our results indicated that pupil responses on delayed trials showed distinct markers of learning that were not present on immediate trials. DDM parameters from delayed trials also showed expected trends for learning. Finally, we demonstrated that DDM parameters elicit differential pupillary responses. Together, these results indicate that learning dynamics and decisional processes can be decoded from pupillary responses.

---

Understanding the dynamic mechanisms underlying categorization, that is, mapping variable sensory signals into discrete categories, is a fundamental pursuit in cognitive neuroscience. There are myriad cognitive processes during category learning that have been related to different neuromodulatory systems that operate on multiple timescales. Pupillometry has emerged as a vital tool for understanding these learning dynamics, as it can offer meaningful insights into the online neurocognitive processes that are involved and the neuromodulatory systems that mediate performance (Beatty, 1982; Foroughi et al., 2017; McHaney et al., 2021, 2023; Reimer et al., 2016; Sibley et al., 2011). In the behavioral domain, drift-diffusion models (DDM) have provided a robust mathematical framework to dissect the perceptual and decisional processes involved in stimulus encoding and response selection (Donkin & Brown, 2018; Ratcliff & McKoon, 2008). Recent computational cognitive neuroscience-focused approaches have applied DDMs to trial outcomes and response times from learning tasks to capture the dynamics of auditory learning (Roark et al., 2021). Therefore, measuring pupillary responses in combination with DDMs may help uncover some of the neural mechanisms driving decision-making during learning (de Gee et al., 2020; Preuschoff et al., 2011).

Relating pupillary responses to DDM parameters requires crucial methodological considerations. Evidence from studies in animals (McGinley et al., 2015; Nelson & Mooney, 2016) and humans (Yokoi & Weiler, 2022; but see Einhäuser et al., 2008) suggest that pupil dilations are affected by motor movements. For example, pupil dilations increase during walking movements in mice (McGinley et al., 2015; Nelson & Mooney, 2016; Vinck et al., 2015) and during motor-driven response selection in humans (Yokoi & Weiler, 2022). However, the role of motor movements on pupillary responses is rarely acknowledged in human studies. Some human pupillometry studies have strategically implemented timing delays in the experimental design to disambiguate motor-related influences from a response in the pupil recordings (e.g., McHaney et al., 2021, 2023; Winn et al., 2015). Yet, delaying trial events to optimize pupillary responses also introduces challenges in analyzing behavioral responses.

When timing delays are introduced in a decision-making task to optimize measurement of the pupil response, response times are confounded because the participant is instructed to withhold their response selection for a given amount of time. DDMs rely on trial outcomes and *precise response times* to derive metrics of decision-making (Bogacz et al., 2010; Donkin & Brown, 2018; Parés-Pujolràs et al., 2021; Ratcliff et al., 2016; Ratcliff & McKoon, 2008). Diffusion models assume that the brain accumulates sensory evidence from a stimulus to make a categorical decision (Ratcliff et al., 2016). This process of evidence accumulation is reflected by an increase in local neuronal firing rates associated with each category alternative (Gold & Shadlen, 2007). When neuronal firing rates associated with a particular category alternative crosses its associated threshold, a categorical decision is made (Brody & Hanks, 2016; Gold & Shadlen, 2007). Instructing participants to delay their physical response selection, which inflates response times, may confound DDM estimation because it is plausible that a category decision has already been determined during the response selection delay. Therefore, DDM parameters calculated from response times when a delay was present may not reflect the decision-making process. This begs the question as to the extent to which DDM parameters calculated with delayed responses to optimize measurement of the pupil response can provide comparable insights into learning dynamics as those from immediate responses, which are typical in a standard learning task.

Delays to optimize pupil measurement may impact the learning process. Learning to map highly variable auditory stimuli to discrete categories can be difficult but can be learned within a few hundred trials using trial-by-trial feedback (Roark et al., 2021; Roark & Chandrasekaran, 2023; Yi & Chandrasekaran, 2016). Prior neuroimaging studies have demonstrated that a cortico- striatal network, including the auditory cortex, prefrontal cortex, and the striatum, is involved during learning (Feng et al., 2018; Yi et al., 2016). This cortico-striatal network helps map auditory stimuli to categories based on non-verbalizable rules that can be fine-tuned by trial-by-trial feedback. Typically, auditory category learning paradigms present trial events in an immediate, sequential manner, wherein there are minimal delays (< ∼500 ms) between stimulus presentation and response selection and between response selection and feedback (e.g., Maddox et al., 2003; Roark & Chandrasekaran, 2023; Yi & Chandrasekaran, 2016). Delaying feedback presentation by even 1000 ms can have detrimental effects on learning performance when the category structures require complex integration across multiple dimensions (Chandrasekaran et al., 2014; Maddox et al., 2003). It is hypothesized that the cortico-striatal network that reinforces learning is disrupted by feedback delays when category structures are partitioned in a complex manner, wherein a single acoustic dimension cannot uniquely identify a category (Chandrasekaran et al., 2014).

In contrast, learning performance is not impacted by delayed feedback presentation when category structures can be simply defined by verbalizable rules (Chandrasekaran et al., 2014; Maddox et al., 2003). Learning categories with a rule-based distinction engages a different set of neural circuits than those that are engaged when category structures involve integration across dimensions. The neural circuits engaged for learning rule-based category structures involve the prefrontal cortex for selective attention and working memory resources (Ashby et al., 1998; Nomura et al., 2006) and the caudate nucleus for feedback processing to fine-tune category rules (Filoteo et al., 2005; Tricomi & Fiez, 2008). While delays in feedback do not impact learning for rule-based category structures, delays between stimulus presentation and response selection may heighten demands on working memory to hold onto stimulus information before a response can be initiated. Working memory is also known to impact pupillary responses (McHaney et al., 2021; Miller et al., 2019; Robison & Unsworth, 2019; Unsworth & Robison, 2017). Therefore, it remains unclear whether delaying trial events to optimize pupil measurement can provide similar insights into learning dynamics.

In the current study, we measured pupillary responses during rule-based auditory category learning and associated the pupillary responses with DDM parameter to further understand decisional processes underlying auditory category learning. The rule-based category structure was chosen to avoid confounds in learning from delayed response feedback. Inherent to our study design, we manipulated the timing of trial events, wherein half of the trials were presented in an immediate manner that is standard for category learning tasks, and the other half of trials had timing delays (2-4 s) between each trial event to optimize pupil measurement with minimal impacts from motor movement. Importantly, we assayed pupillary responses between delayed and immediate trials through several lenses to understand the potential impacts of timing delays.

First, we examined the extent to which pupillary signatures of learning were preserved in immediate trials, with more motor contaminations, relative to delayed trials. Prior work has demonstrated greater differentiation between pupillary responses time-locked to stimulus presentation on correct versus incorrect trials, prior to categorization and feedback, with learning progress (McHaney et al., 2021), as well as an overall decrease in pupil size with learning progress (Beatty, 1982; Foroughi et al., 2017; McHaney et al., 2021; Sibley et al., 2011). As such, we hypothesized that these pupillary signatures of learning would be less pronounced on immediate trials due to motor contaminations, compared to delayed trials.

Furthermore, we examined the extent to which accuracies and response times were preserved in delayed trials in a manner that could allow the use of DDMs. If accuracies and response times were highly correlated across immediate and delayed trials, then meaningful DDM parameters could be estimated from delayed trials. Prior research applying DDMs to rule-based auditory category learning found that evidence accumulation rates remained stable across learning, while decision thresholds decreased with learning (Roark et al., 2021). If the DDMs calculated from delayed trials in our study are equivalent to those from a standard rule-based learning task, then we should also observe stable evidence accumulation rates and a decrease in decision thresholds across learning. The ability to extract meaningful DDM parameters from delayed trials is critical to understanding how stimulus processing, as indexed by pupillary responses, impacts decisional processes during learning.

## Methods

### Participants

Eighteen adults (16 females; 2 males) between the ages of 18 – 26 (*M* = 20.72, *SD* = 1.90) participated in this study. Participants were recruited from the University of Pittsburgh’s Department of Communication Science and Disorders research participant pool and received course credit for participation in this study. All participants were native speakers of English and were required to have air conduction thresholds ≤ 25 dB at 250, 500, 1000, 2000, 4000, and 8000 Hz. This research protocol was approved by the Institutional Review Board at the University of Pittsburgh.

### Stimuli

Stimuli were complex nonspeech ripples that varied in spectral and temporal modulation. The category distributions were created by randomly sampling from a two-dimensional Gaussian distribution and then mirroring that distribution across an optimal category boundary along a temporal modulation dimension. As such, the two categories had identical distributions and could be optimally distinguished from one another by a rule along the temporal modulation dimension (Fig. 1A). Hence, these stimuli were divided into rule-based auditory categories. The categories were somewhat probabilistic in that there were a few stimuli in each category that fell on the opposite side of a category boundary, making them an outlier of their true category or an exception to the rule. The stimuli were generated using custom scripts in MATLAB (Mathworks Inc., Natick, MA) and were amplitude matched at 70 dB.

**Figure 1.**
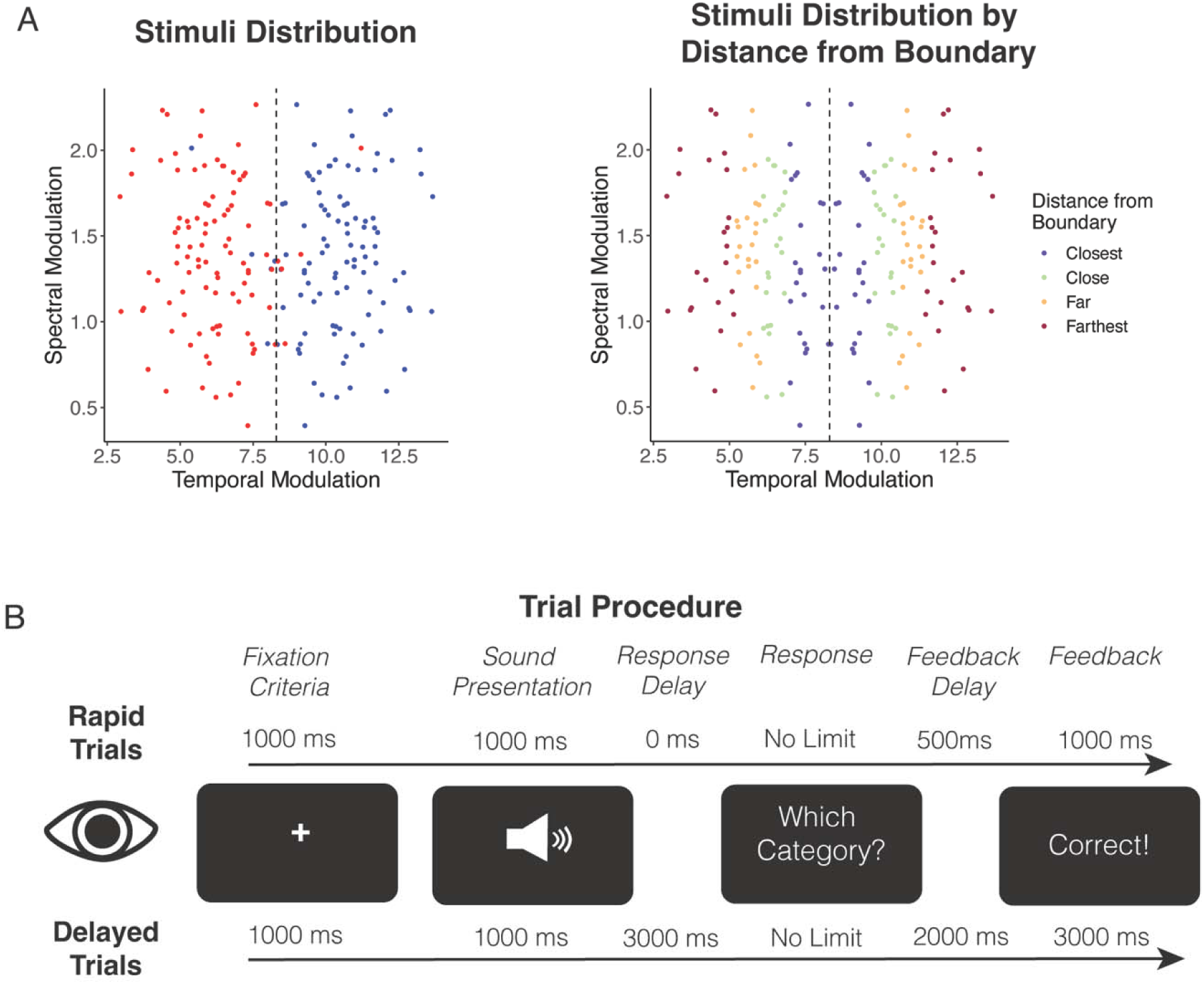
Stimuli and Procedure. A) Left panel: Stimuli distribution across the spectral and temporal modulations color-coded by category. Vertical dashed line indicates category boundary. Right panel: Stimuli distribution color-coded by distance from the categorical boundary. B) Single-trial procedure with trial-event timings for immediate trials and delayed trials in order by fixation cross, sound presentation, delay between sound presentation and response prompt, unlimited response time, delay between response and feedback, and feedback display.

### Procedure

Pupillometry was recorded while participants completed a category learning task. Participants were seated with their head stabilized on a chin and forehead rest during recording. Luminance of the visual field was controlled across participants by maintaining consistent dim lighting in the testing room. Left-eye pupil sizes were recorded at 1000 Hz with an Eyelink 1000 Plus Desktop Mount. A nine-point eye tracking calibration was performed prior to beginning the category learning task.

Participants learned to categorize auditory stimuli across four blocks of 50 trials, for a total of 200 trials. Half of the trials (n = 100) were ‘immediate’ trials, wherein trial events were timed for a standard category learning paradigm. The other half of trials were ‘delayed’ trials, wherein there was a 2-4 second delay between each trial event. The delayed trials were included in order to increase the interpretability of the pupil response data. Delayed and immediate trials were presented in random order throughout the task. Each trial began with a fixation cross that was presented on the center of the screen. The cross was color coordinated to indicate the trial type to the participant. A green cross indicated immediate trials and a red cross indicated delayed trials. Participants were required to look at the cross for a minimum of 1000 ms to trigger the start of the auditory stimulus. The timing of the remaining events in the trial differed depending on the type of trial.

On immediate trials, the auditory stimulus was presented for 440 ms and participants were immediately prompted to respond with the category via button press. Participants had no time limit to make their category response. After the button press, there was a 500 ms delay before corrective feedback was displayed on the screen in the form of ‘Correct’ or ‘Wrong’. Feedback was displayed for 1000 ms. On delayed trials, there was a 4000 ms delay after the stimulus was presented to allow changes in pupil size in response to the auditory stimulus before the category response prompt was presented on the screen. Participants had unlimited time to make their category response via button press. There was a 2000 ms delay after the button press before the corrective feedback was displayed on the screen for 3000 ms. Example trials are depicted in Figure 1B.

Participants were provided breaks between each block. Manual drift-correction was performed at the start of each successive block to account for slight drifts in pupil tracking, which helped maintain high-quality tracking of the pupil throughout the task. Experiment presentation was controlled using MATLAB version 2018b (Mathworks, Inc.).

### Category Learning Data Analysis

Accuracy and response time data from the category learning task were analyzed using R version 4.3.1 (R Core Team, 2023). We used the *lme4* package (Bates et al., 2015) to analyze accuracies on immediate versus delayed trials in a binomial generalized linear mixed effects model. The outcome variable of the model was trial-by-trial outcome (correct, incorrect). The model included fixed effects of trial number, trial type (reference = delayed), and the interaction of trial number and trial type. The maximal random effect structure of the model that promoted convergence and a non-singular fit contained a random slope of trial type per participant and a random intercept of stimulus token: *trial Outcome ∼ trial number * trial type + (trial type | participant) + (1 | stimulus token)*. A similar linear mixed effects model was fit to compare trial-by-trial response times between immediate and delayed trials with the same fixed effects and random effects structure as the binomial generalized linear mixed effects model. Here, response times were calculated as the time taken to provide a button response in reference to the ‘Which Category’ prompt on screen.

We also examined accuracy based on stimuli distance from the optimal category boundary as a proxy for difficulty of decision-making. Stimuli closer to the boundary were considered to be more difficult to categorize, while those farthest from the boundary were considered to be easier to categorize. Trials for the outlier/exception stimuli (e.g., category A tokens on the category B side; see Fig 1A) were first removed from this analysis. These stimuli were removed in order to focus on stimuli that were on the correct side of the optimal category boundary. The distance from the optimal boundary was calculated for each stimulus token. The stimuli were then divided into four equal sized groups, referred to as bins from hereon, based on the absolute value of this distance from the optimal category boundary (Fig 1A). Average accuracy per block for each stimulus bin was calculated for each participant. A linear mixed effects model was fit with an outcome variable of block accuracy, fixed effects of trial type (reference = delayed), block number, bin (reference = closest bin), and all two- and three-way interactions between fixed effects. The maximal random effect structure that provided a non-singular fit and promoted convergence comprised of a random slope for trial type per participant and a random intercept for the interaction between participant and block.

### Drift-Diffusion Modeling

We used the Bayesian implementation of the drift-diffusion model (DDM) developed by Paulon et al. (2020) in R version 4.3.1 (R Core Team, 2023) with the *lddmm* package (Paulon & Sarkar, 2023) to analyze trial accuracies and response times from the category learning task. The DDM estimates an evidence accumulation rate parameter *m_d,s_* and a decision threshold parameter *b_d,s_* for each combination of category *s* and response *d*. Evidence accumulation rate reflects the quality of information extracted from the stimulus, with higher evidence accumulation rates reflecting more efficient extraction of information from the stimulus. Decision threshold reflects the amount of information that must be accumulated before a decision is made and is also a metric of response caution (Bogacz et al., 2010). Higher decision thresholds reflect more cautious responding, while lower decision thresholds indicate more impulsive responding. The DDM also fits an offset parameter *d_s_* for each category that reflects the time taken by actions that are not directly relevant to the decision process, such as encoding the stimulus before evidence accumulation starts, and the time taken for the physical action of pressing a button to make the response. Importantly, the DDM allows the *m_d,s_* and *b_d,s_* parameters to vary between participants to control for the substantial variability across participants. The DDM also allows the *m_d,s_* and *b_d,s_* parameters to change slowly over time across learning blocks. This helps to account for the changes in decision-making processes during learning.

We estimated the DDM parameters separately for immediate and delayed trials using a Bayesian framework that assigned priors to the parameters and relied on samples drawn from the posterior using a Markov chain Monte Carlo algorithm for estimation and inference. Prior to estimating the DDM parameters, the top and bottom one percent of trials based on response times were removed as outliers to improve the model fit. Then, the algorithm was run for 6000 iterations with the initial 2000 iterations discarded as burn-in. The remaining iterations were thinned in intervals of 5 to reduce autocorrelation. Consistent with prior research, we focus on the DDM parameters associated with correct responses only because gradual improvements in correct categorical decisions characterizes learning (Roark et al., 2021, 2022). Posterior means are reported as point estimates and 95% pointwise credible intervals were used to assess uncertainty.

### Decision-bound Computational Modeling

We also used decision-bound computational models to understand how participants grouped the stimuli into categories, which informs on the categorization strategies that were used during learning (Ashby, 1992; Ashby & Maddox, 1993; Chandrasekaran et al., 2016; McHaney et al., 2021). These models assume that participants divide the stimuli into categories in a two-dimensional space varying along the temporal modulation and spectral modulation dimensions with decision boundaries that can rely on rule-based or procedural-based learning processes. Decision bound models were originally developed for the visual modality (Ashby & Maddox, 1993) but have commonly been used in the auditory modality, including to examine auditory category learning (Chandrasekaran et al., 2016; Goudbeek et al., 2008, 2009; Maddox et al., 2002, 2013, 2014; Maddox & Chandrasekaran, 2014; McHaney et al., 2021; Roark & Holt, 2019; Scharinger et al., 2013; Smayda et al., 2015). Models are fit individually for each participant and learning block to mitigate challenges of interpreting model fits to aggregate data (Ashby et al., 1994; Estes, 1956; Estes & Maddox, 2005). We fit three classes of models with multiple instantiations within each class separately for delayed trials and immediate trials. These classes included *rule-based models, a general linear classifier (GLC) model, and the inconsistent/random responder model*.

### Rule-based Models

Rule-based models reflect hypothesis-testing mechanisms that are mediated by the reflective learning system. These models assume that participants selectively attend to a single acoustic dimension during learning and draw their categorical boundary along this single dimension. These unidimensional models have two free parameters. One parameter reflects the location of the categorical boundary along the relevant dimension. The other parameter reflects perceptual and criterial noise. We fit separate unidimensional models that assumed participants made categorization decisions using only the relevant spectral dimension or only the temporal dimension.

### General Linear Classifier Model

The GLC model assumes there is a linear decision boundary that assumes that participants integrate both dimensions to divide the categories and that the boundaries are not orthogonal to the dimensions. The GLC model requires fitting of three free parameters. Two parameters reflect the slope and intercept of the boundary. The other parameter reflects decisional and criterial noise.

### Inconsistent/Random Responder Model

The inconsistent/random responder model assumes that participants categorize stimuli with all responses being equally probable. Therefore, this model captures participants who randomly respond to the stimuli or those who change their strategy too often to be captured by either of the other models.

### Model Fitting

Several versions of each model were separately fit for delayed and immediate trials with different assumptions regarding the assignment of responses to the perceptual space to account for differences in how participants responded to stimuli in different regions of the space. This manner of model fitting ensures that the best-fit model is invariant to different stimulus-response region assignments (Ashby, 1992; Ashby & Maddox, 1993). In total, eleven unidimensional, one GLC model, and one random responder model were fit per block for each participant. Models were separately fit for immediate trials and delayed trials by splitting learning trials into early and late blocks for each participant. Given that the learning task contained 100 delayed trials and 100 immediate trials in random order, we considered the first 50 delayed trials as ‘early’ and the remaining 50 delayed trials as ‘late’, with the same considerations taken for the immediate trials. In total, there were 252 models fit per trial type across participants (7 models x 18 participants x 2 blocks = 252 models per trial type).

All model parameters were estimated using maximum likelihood procedures (Wickens, 1982) and model selections were based on the Bayesian Information Criteria (BIC). The BIC penalizes models with more free parameters: BIC = r * ln(N) - 2 ln(L), where *r* is the number of free parameters, *N* is the number of trials in a given block for each participant, and *L* is the likelihood of the model based on the data (Schwarz, 1978). For each participant and block, the model among all thirteen individual models across each model class with the lowest BIC was selected as best-fit (McHaney et al., 2021).

### Assessment of Model Fit

We separately computed the proportion of participant’s responses accurately predicted by the best fit model for delayed and immediate trials as a measure of model fit to understand how well the best-fit model for each participant and block accounted for each participant’s actual response pattern based on the given trial type. We calculated a *predicted* response based on the fitted parameters from the best-fit model for each participant and block. The predicted response reflected the category response (i.e., A or B) that the best-fit model predicted the participant would have made if they applied the selected strategy throughout that entire block. We then compared the predicted response to the *observed* response made by the participant for each stimulus in each block as a measure of model accuracy. When the predicted response matched the observed response, it was coded as a ‘Correct’ prediction, while predicted responses that did not match the observed response were coded as ‘Incorrect”. This model accuracy measure served as a metric of goodness-of-fit.

We found that the best-fit models accurately captured participants’ response patterns in the early half 76.8% of the time for delayed trials and 74.5% of the time for immediate trials. In the late half, model accuracy was 79.3% for delayed trials and 81.9% for immediate trials. The chance-level of success was 50% for a single trial and for each trial type. Therefore, the model fits were greater than chance, indicating that the best-fit models provided an accurate account of participants’ response patterns.

### Pupillary Data Processing

Raw pupillometry data were processed in R version 4.3.1 (R Core Team, 2023) using the *eyelinker* package (Barthelme, 2023). Data were first down sampled to a sampling rate of 50 Hz, and then were processed to remove noise, such as blinks and saccades. Trials where more than 20% of the samples were missing due to blinks or saccades (*n* = 212 out of 3600 trials) were excluded from analysis (Winn et al., 2018). Remaining trials that contained missing samples due to blinks or saccades were linearly interpolated from 120 ms prior to the onset of the blink or saccade to 120 ms after the blink or saccade (McHaney et al., 2023). The pupillary response data were then baseline corrected and normalized on a trial-by-trial basis using the average pupil size across the 500 ms baseline period that immediately preceded 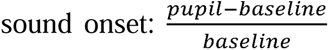. This baseline correction and normalization method was performed to account for a downward drift in baseline that tends to occur across a task and for individual differences in pupil dynamic range (Winn et al., 2018). The resulting pupillary data were reported as the percent change in pupil size relative to baseline.

### Pupillary Data Analysis

To examine pupillary responses time-locked to stimulus onset, we calculated growth curve analyses (GCA; Mirman, 2014). GCA uses orthogonal polynomials to capture distinct functional forms of the pupillary response to examine how the pupillary response unfolded overtime. We fit a fourth-order polynomial GCA to model the pupillary response from 0 – 3000 ms time-locked to the onset of stimulus presentation for both immediate and delayed trials. This time period was chosen in an attempt to avoid pupil changes caused by preparation for motor execution on delayed trials that may occur closer to the 4000 ms point when a response is initiated (de Gee et al., 2014; Murphy et al., 2016; Vinck et al., 2015). This fourth-order GCA fit five parameters to map the shape of the pupillary response. The intercept reflects the overall change in the pupillary response over the entire time window and can be interpreted as the mean change in pupil size over time. The linear (ot1) term represents the slope of the pupillary response. If the linear term is positive, this reflects the rate of dilation, while a negative ot1 term reflects the rate of constriction. The quadratic (ot2) term reflects the curvature of the pupillary response, wherein a larger, negative quadratic term suggests a more parabolic shape and values closer to zero reflect a more linear shape. The cubic (ot3) and quartic (ot4) terms reflect the extent to which secondary and tertiary inflection points occur in the pupil response, respectively (Kuchinsky et al., 2014; McGarrigle et al., 2017). The absolute value of each term reflects the strength of the pupil response, while the polarity of the term reflects the direction of the response. All GCAs were conducted using the *lme4* package (Bates et al., 2015) with log-likelihood maximization using the BOBYQA optimizer to promote convergence. The *lmerTest* package (Kuznetsova et al., 2017) was used to estimate *p*-values.

We first estimated a GCA to examine the effect of trial type on trials that were categorized correctly and incorrectly. The model contained fixed effects of each growth curve term, trial type, trial outcome, and all two- and three-way interactions between growth curve terms, trial type, and trial outcome. The maximal model that promoted convergence and a non-singular fit had a random effect structure containing a random slope of each growth curve term per participant, a random intercept of the interaction of participant and trial type, a random intercept for the interaction of participant and trial outcome, and a random intercept for the three-way interaction between participant, trial outcome, and trial type.

Next, we fit a GCA to examine the extent to which pupillary responses decreased with learning. This model contained fixed effects of each growth curve term, trial type, learning block, and all two- and three-way interactions between fixed effects. The maximal random effect structure that promoted convergence and a non-singular fit consisted of a random slope of each growth curve term per participant and a random intercept for the interaction of participant and trial type.

As a metric of difficulty, we estimated a GCA of pupillary responses on each trial type based on the stimulus distance from boundary. This model included fixed effects of each growth curve term, trial type, stimulus distance to boundary, and all two- and three-way interactions between growth curve terms, trial type, and stimulus distance to boundary. The maximal model that promoted convergence and a non-singular fit had a random effect structure comprised of a random slope of each growth curve term per participant and a random intercept of the interaction of participant and trial type.

We also estimated separate GCAs to examine the extent to which pupillary responses reflected DDM parameters of decision-making on correct trials. The models included fixed effects of each growth curve term, trial type, DDM parameter (either evidence accumulation rates or decision thresholds), and all two- and three-way interactions between growth curve term, trial type, and DDM parameter. For the evidence accumulation rate GCA, the maximal model that promoted convergence and a non-singular fit included a random slope of each growth curve term per participant, a random intercept of the interaction of participant and trial type, a random intercept for the interaction of participant and evidence accumulation rate, and a random intercept for the three-way interaction between participant, evidence accumulation rate, and trial type. For the decision threshold GCA, the maximal model included a random slope of each growth curve term per participant and a random intercept of the interaction of participant and trial type.

## Results

### Pupillary Responses

#### Baseline Pupil Size Does Not Differ by Trial Type

We first examined baseline pupillary responses for the 500 ms prior to stimulus onset between immediate and delayed trials to understand whether pre-trial arousal levels differed on delayed versus immediate trials (Fig. 2A). The baseline pupil size was calculated separately for immediate and delayed trials as the average pupil size from -500 to 0 ms time-locked to stimulus onset, prior to the baseline normalization step described in the *Pupillary Data Processing* section. Baseline pupil sizes are reported in arbitrary units. The distributions of baseline pupil sizes were non-normal, based on visual inspection and significant Shapiro-Wilks test. Therefore, a Wilcoxon matched pairs signed rank test was performed to examine differences in baseline pupil sizes as a function of trial. The results revealed that the median average baseline pupil size for delayed trials (*Mdn* = 1363.7) was not different from immediate trials (*Mdn* = 1378.0; *p* = .495, *r* = .169, 95% CI = [-34.798,16.974]). In summary, pre-trial arousal levels as measured by baseline pupil size were not different between trial types.

**Figure 2.**
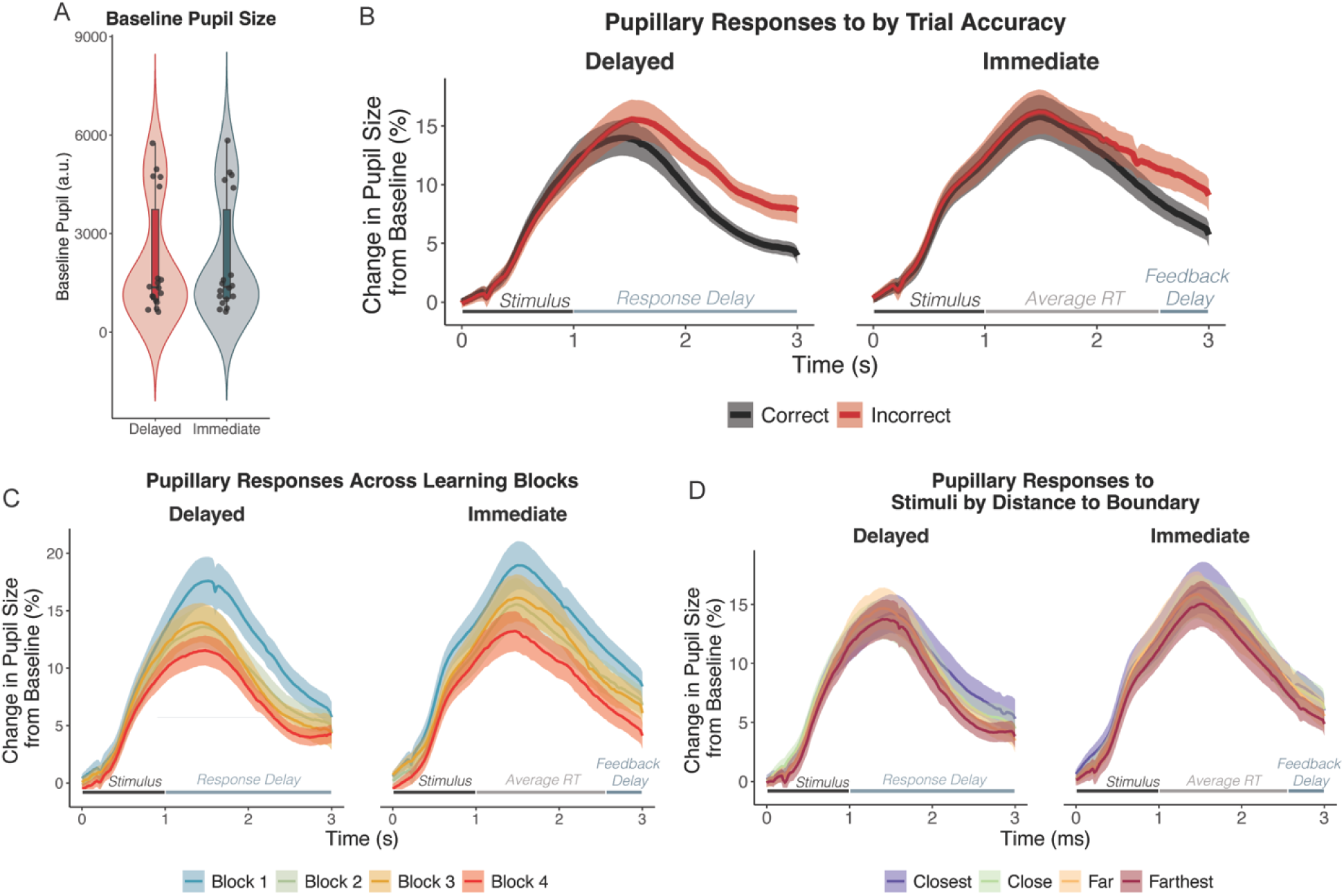
Pupillary responses between immediate and delayed trials. A) Distribution of the average baseline pupil size in arbitrary units (a.u.) for the 500 ms preceding stimulus onset for immediate and delayed trials. B) Pupillary responses for correct (black) and incorrect (red) trials. C) Pupillary responses for each learning block on delayed and immediate trials showing dynamic changes in the pupil response with learning task progression D) Pupillary responses for stimuli based on distance from the categorical boundary. Responses to stimuli closest to the categorical boundary are more difficult to categorize than those farthest from the boundary. B-D) Solid lines denote the mean pupillary responses, and the shaded regions denote standard error of the mean. Lines near the X-axis represent average timing of trial events. RT = response time.

### Pupillary Responses on Delayed Trials Better Reflect Trial Outcome

We used GCAs to analyze the pupillary response 0-3000 ms time-locked to the onset of the auditory stimulus on each trial to examine the extent to which the type of trial (immediate vs. delayed) and trial outcome (correct vs. incorrect) impacted the pupil response (Fig. 2B). Crucially, the 0-3000 ms time-period encompassed different trial events, based on the type of trial. On delayed trials, the 0-3000 ms time-period measured changes in pupil size in response to the auditory stimulus while waiting to make the physical categorical button press. On immediate trials, the time-period of interest, on average, encompassed auditory stimulus presentation, categorical button press, and the 500 ms delay prior to feedback presentation.

First, we examined the effects of trial type on correct trials (Table 1). We observed main effects of the intercept, linear (ot1), quadratic (ot2), cubic (ot3), and quartic (ot4) time terms between immediate and delayed trials on trials with a correct categorization choice. Specifically, immediate trials had a larger overall pupillary response, a steeper slope, a stronger primary inflection point, and flatter secondary and tertiary inflection points compared to pupillary responses on delayed trials. Thus, pupil dilation on immediate trials with correct categorization were larger and dynamically differed from those on delayed trials.

**Table 1.** Fixed-effect estimates for model of pupillary responses from 0 to 3000 ms time-locked to stimulus onset to examine the effect of trial type and trial outcome (observations = 10,872, groups: participant x Outcome x Trial Type = 72; participant x Outcome = 36, participant x Trial Type = 36, participant = 18).

| Fixed Effect | Estimate | SE | 95% CI | *t* | *p* |
| --- | --- | --- | --- | --- | --- |
| Intercept | 8.01 | 1.01 | [6.04, 9.98] | 7.96 | < .001 |
| ot1 | 7.71 | 3.75 | [0.37, 15.06] | 2.06 | 0.040 |
| ot2 | -48.65 | 5.79 | [-60.01, -37.29] | -8.40 | < .001 |
| ot3 | 10.58 | 2.22 | [6.23, 14.92] | 4.77 | < .001 |
| ot4 | 14.07 | 2.23 | [9.70, 18.44] | 6.31 | < .001 |
| Trial Type (Delayed vs. Immediate) | 1.73 | 0.56 | [0.64, 2.83] | 3.10 | 0.002 |
| ot1 x Trial Type | 11.68 | 0.55 | [10.60, 12.75] | 21.36 | < .001 |
| ot2 x Trial Type | -1.37 | 0.55 | [-2.44, -0.30] | -2.51 | 0.012 |
| ot3 x Trial Type | -6.07 | 0.55 | [-7.14, -4.99] | -11.10 | < .001 |
| ot4 x Trial Type | -3.13 | 0.55 | [-4.20, -2.06] | -5.73 | < .001 |
| Outcome (Correct vs. Incorrect) | 1.58 | 0.52 | [0.57, 2.59] | 3.07 | 0.002 |
| ot1 x Outcome | 19.49 | 0.55 | [18.42, 20.56] | 35.66 | < .001 |
| ot2 x Outcome | 1.21 | 0.55 | [0.14, 2.28] | 2.22 | 0.027 |
| ot3 x Outcome | -6.99 | 0.55 | [-8.06, -5.92] | -12.79 | < .001 |
| ot4 x Outcome | -0.38 | 0.55 | [-1.45, 0.69] | -0.69 | 0.490 |
| Outcome x Trial Type | -0.51 | 0.59 | [-1.67, 0.66] | -0.85 | 0.394 |
| ot1 x Outcome x Trial Type | -7.04 | 0.77 | [-8.55, -5.52] | -9.11 | < .001 |
| ot2 x Outcome x Trial Type | 4.44 | 0.77 | [2.93, 5.96] | 5.75 | < .001 |
| ot3 x Outcome x Trial Type | 7.70 | 0.77 | [6.18, 9.21] | 9.96 | < .001 |
| ot4 x Outcome x Trial Type | -1.59 | 0.77 | [-3.11, -0.08] | -2.06 | 0.039 |

Growth curve formula: $\text{lmer}(\text{Pupil} \sim (\text{ot1} + \text{ot2} + \text{ot3} + \text{ot4}) * \text{Outcome} * \text{Trial Type} + (\text{ot1} + \text{ot2} + \text{ot3} + \text{ot4} / \text{participant}) + (1 / \text{participant} : \text{Trial Type}) + (1 / \text{participant} : \text{Outcome}) + (1 / \text{participant} : \text{Outcome} : \text{Trial Type}), \text{control} = \text{lmerControl}(\text{optimizer} = \text{"bobyqa"})$ . Orthogonal polynomial terms: ot1 = linear; ot2 = quadratic; ot3 = cubic; ot4 = quartic.

Next, we examined the extent to which trial type modulated pupillary responses on incorrect trials, relative to correct trials. For incorrect trials, there were significant effects of trial type on the intercept, linear, quadratic, and cubic time terms for incorrect trials. This indicates that pupillary responses on delayed trials were larger overall (intercept), faster to dilate overtime (ot1), and had flatter curvatures of the primary (ot2) and secondary (ot3) inflection points when the trial was categorized incorrectly, compared to trials with correct categorization. We also observed significant three-way interactions between trial outcome and trial type on the linear, quadratic, cubic, and quartic time terms. These interactions indicate that there were smaller differences in the overall size of the pupil response (intercept), the rate of dilation (ot1), and the primary (ot2), secondary (ot3), and tertiary (ot4) inflection points between correct and incorrect responses on immediate trials, relative to delayed trials. These findings demonstrate that delaying trial events resulted in more robust, dynamic differences between correct and incorrect trials, compared to immediate timing as in a traditional learning paradigm format. Additionally, the differences observed between correct and incorrect trials for immediate trials may arise from the combination of sensory, decisional, and motor processes because the interest period encompassed several trial events. Therefore, pupillary responses on delayed trials provided more meaningful output associated with behavior than immediate trials.

### Pupil Response Changes Across Learning are More Apparent on Delayed Trials

We examined average pupillary responses across learning blocks to assess effects of learning between immediate and delayed trials (Fig. 2C). We observed a significant effect of Block on all time terms for delayed trials (Table 2). This suggests that the overall change and the rate of dilation decreased, the primary inflection point became flatter, and the secondary and tertiary inflection points became sharper with progressive learning blocks. Compared to delayed trials, pupillary responses on immediate trials had a smaller change in the primary and secondary curvatures across learning blocks. Additionally, we observed a significant effect of trial type on the intercept, linear, quadratic, and quartic time terms in block 1. These significant effects indicate that immediate trials had a larger overall change, a faster rate of dilation, and a flatter primary and tertiary curvature compared to delayed trials in block 1 at the beginning of learning, which again may arise from the combination of sensory, decisional, and motor processes occurring during the period of interest. Collectively, these findings demonstrate that pupillary responses decreased with task progression, but responses on delayed trials showed greater changes than immediate trials, particularly in the primary and secondary inflection points. Thus, the ability to detect effects of learning from the pupil response was more robust on delayed trials than immediate trials.

**Table 2.**
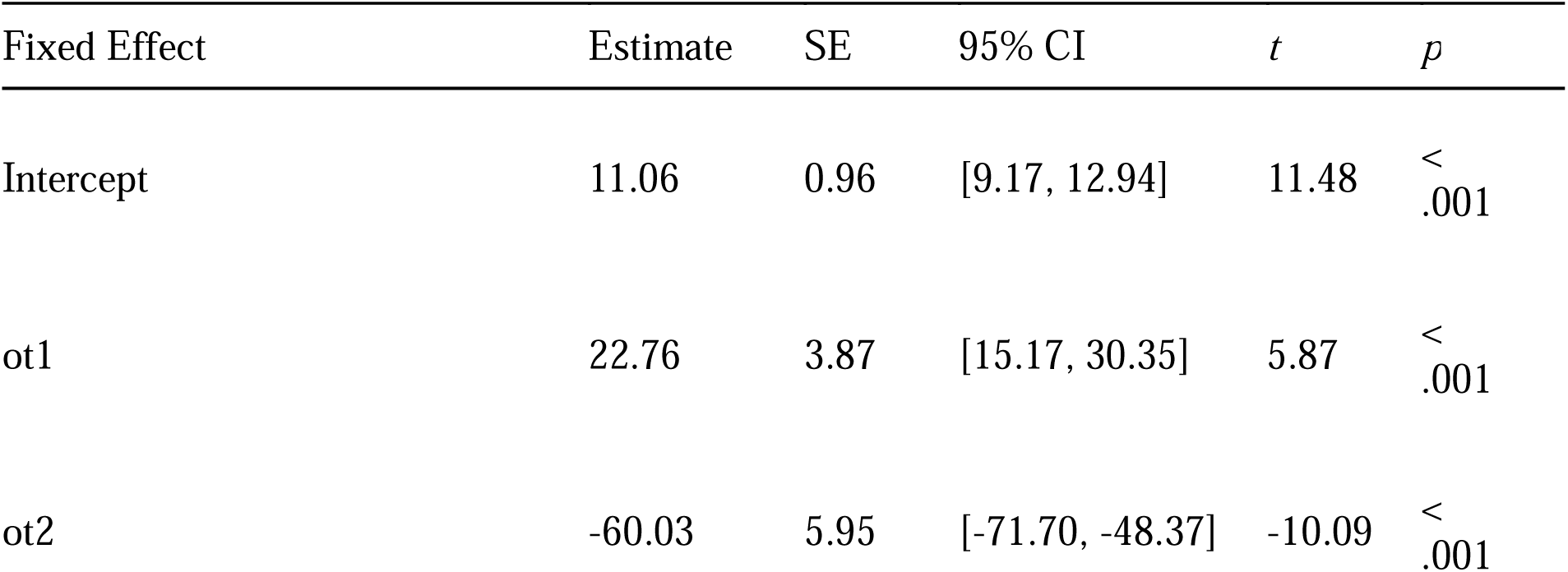
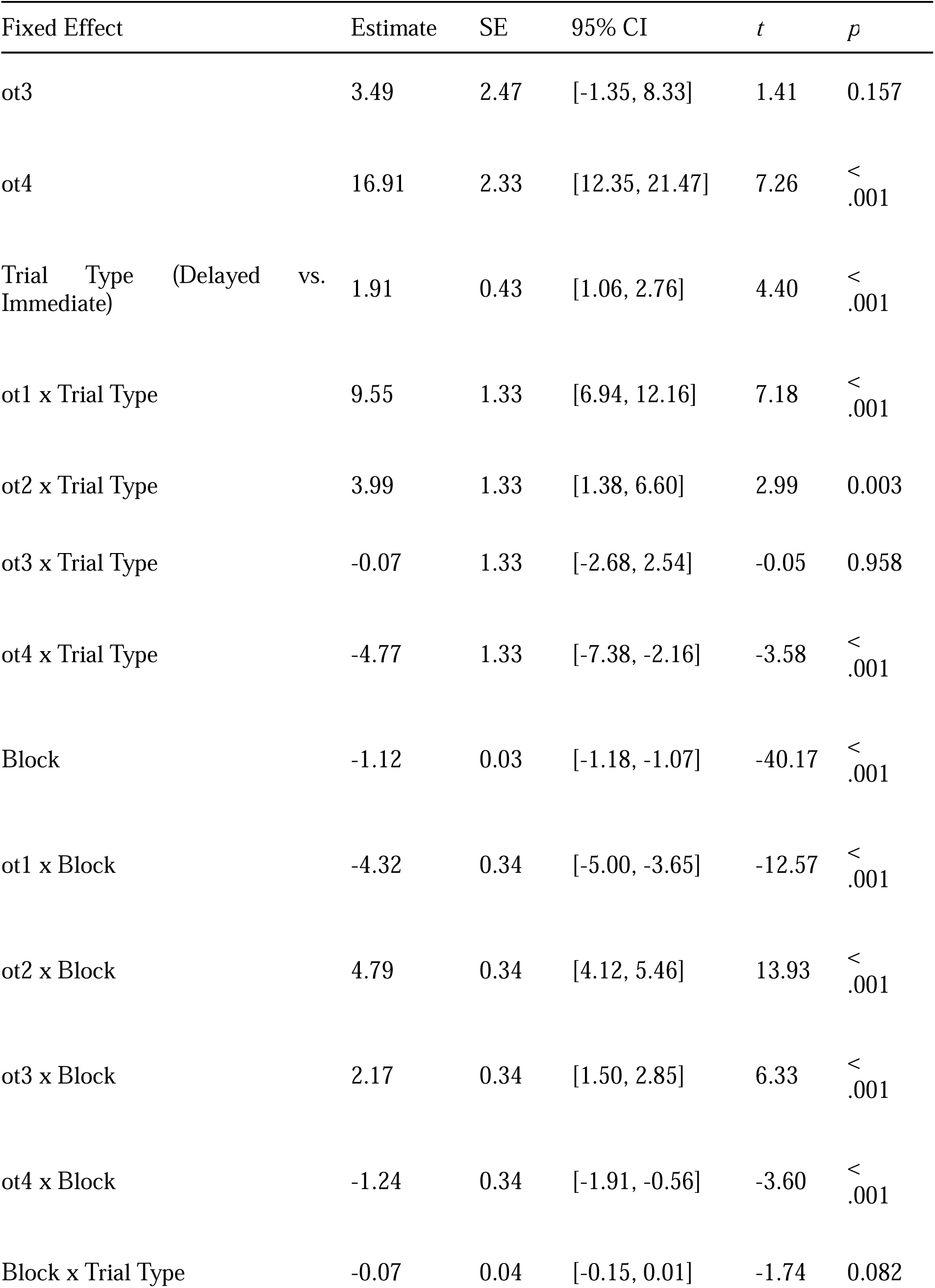
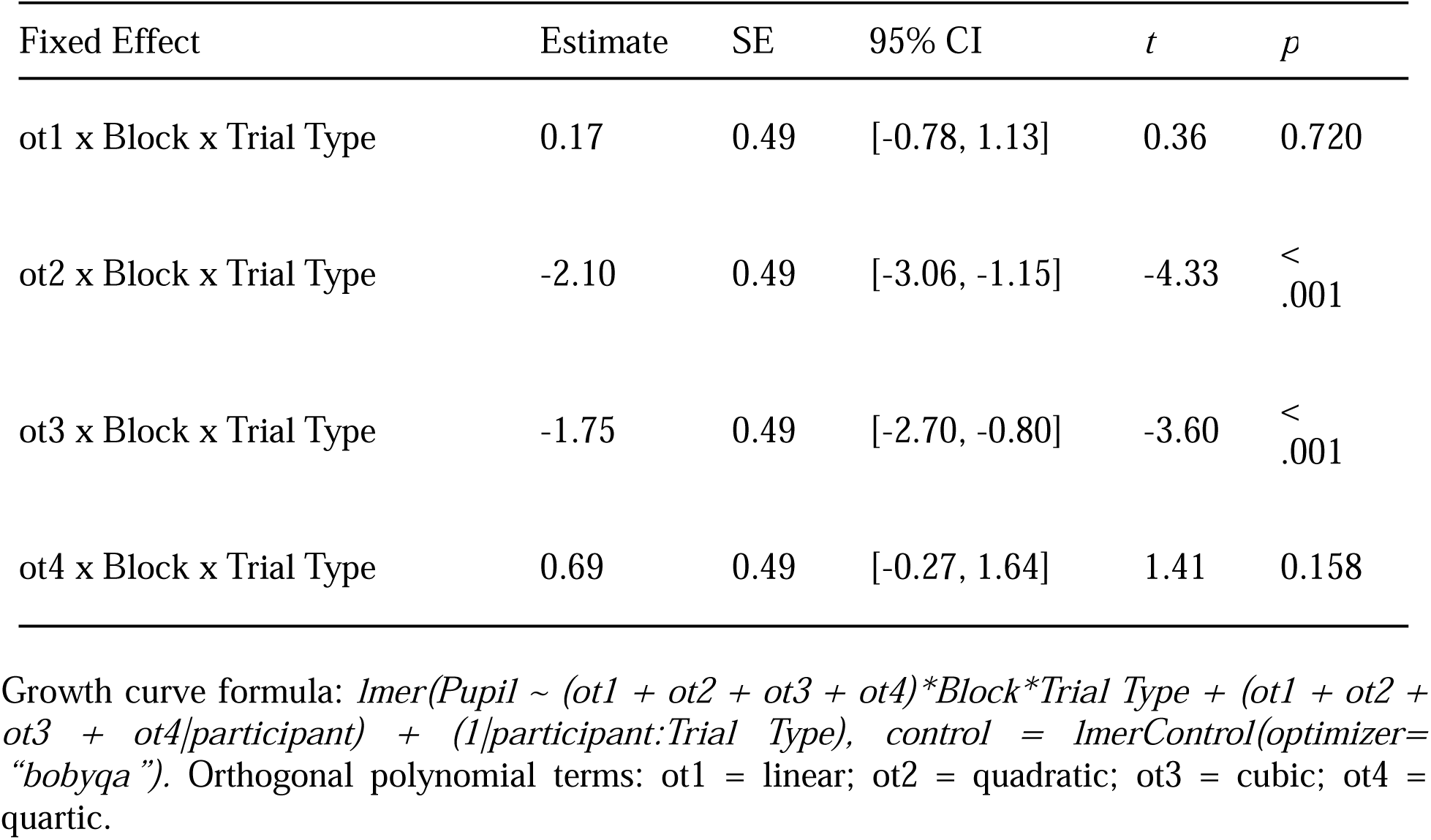
Fixed-effect estimates for model of pupillary responses from 0 to 3000 ms time-locked to stimulus onset to examine the effect of trial type and learning block (observations = 21,744, groups: participant x Trial Type = 36, participant = 18).

| Fixed Effect | Estimate | SE | 95% CI | *t* | *p* |
| --- | --- | --- | --- | --- | --- |
| Intercept | 11.06 | 0.96 | [9.17, 12.94] | 11.48 | < .001 |
| ot1 | 22.76 | 3.87 | [15.17, 30.35] | 5.87 | < .001 |
| ot2 | -60.03 | 5.95 | [-71.70, -48.37] | -10.09 | < .001 |
| ot3 | 3.49 | 2.47 | [-1.35, 8.33] | 1.41 | 0.157 |
| ot4 | 16.91 | 2.33 | [12.35, 21.47] | 7.26 | <.001 |
| Trial Type (Delayed vs. Immediate) | 1.91 | 0.43 | [1.06, 2.76] | 4.40 | <.001 |
| ot1 x Trial Type | 9.55 | 1.33 | [6.94, 12.16] | 7.18 | <.001 |
| ot2 x Trial Type | 3.99 | 1.33 | [1.38, 6.60] | 2.99 | 0.003 |
| ot3 x Trial Type | -0.07 | 1.33 | [-2.68, 2.54] | -0.05 | 0.958 |
| ot4 x Trial Type | -4.77 | 1.33 | [-7.38, -2.16] | -3.58 | <.001 |
| Block | -1.12 | 0.03 | [-1.18, -1.07] | -40.17 | <.001 |
| ot1 x Block | -4.32 | 0.34 | [-5.00, -3.65] | -12.57 | <.001 |
| ot2 x Block | 4.79 | 0.34 | [4.12, 5.46] | 13.93 | <.001 |
| ot3 x Block | 2.17 | 0.34 | [1.50, 2.85] | 6.33 | <.001 |
| ot4 x Block | -1.24 | 0.34 | [-1.91, -0.56] | -3.60 | <.001 |
| Block x Trial Type | -0.07 | 0.04 | [-0.15, 0.01] | -1.74 | 0.082 |
| ot1 x Block x Trial Type | 0.17 | 0.49 | [-0.78, 1.13] | 0.36 | 0.720 |
| ot2 x Block x Trial Type | -2.10 | 0.49 | [-3.06, -1.15] | -4.33 | < .001 |
| ot3 x Block x Trial Type | -1.75 | 0.49 | [-2.70, -0.80] | -3.60 | < .001 |
| ot4 x Block x Trial Type | 0.69 | 0.49 | [-0.27, 1.64] | 1.41 | 0.158 |

Growth curve formula: *lmer*(*Pupil* ~ (*ot1* + *ot2* + *ot3* + *ot4*)\**Block*\**Trial Type* + (*ot1* + *ot2* + *ot3* + *ot4*)/*participant*) + (1/*participant*:*Trial Type*), *control* = *lmerControl*(*optimizer*=“*bobyqa*”). Orthogonal polynomial terms: *ot1* = linear; *ot2* = quadratic; *ot3* = cubic; *ot4* = quartic.

### Pupil Responses on Delayed Trials Reflect Stimulus Difficulty

We also examined differences in the pupillary response by trial type and *stimulus difficulty*, which was defined by four equal-sized groups based on the distance of the stimulus from the optimal category boundary (Fig. 2D; Table 3). For immediate trials, increasing distance from category boundary resulted in less robust changes in the pupillary response, relative to delayed trials. Specifically, the pupillary changes with increasing distance from boundary on the linear, quadratic, cubic, and quartic time terms on immediate trials was significantly smaller than for delayed trials. These results demonstrate that pupillary responses on delayed trials better captured stimulus difficulty compared to immediate trials.

**Table 3.** Fixed-effect estimates for model of pupillary responses from 0 to 3000 ms time-locked to stimulus onset to examine the effect of stimulus difficulty as defined by distance from category boundary (observations = 21,744, groups xparticipant x TrialType = 36, participant = 18).

| Fixed Effect | Estimate | SE | 95% CI | *t* | *p* |
| --- | --- | --- | --- | --- | --- |
| Intercept | 9.11 | 0.94 | [7.27, 10.95] | 9.71 | < .001 |
| ot1 | 14.78 | 4.02 | [6.91, 22.66] | 3.68 | < .001 |
| ot2 | -45.20 | 5.96 | [-56.88, -33.53] | -7.59 | < .001 |
| ot3 | 7.84 | 2.61 | [2.72, 12.95] | 3.00 | 0.003 |
| ot4 | 9.01 | 2.22 | [4.65, 13.37] | 4.05 | < .001 |
| Trial Type (Delayed vs. Immediate) | 1.61 | 0.43 | [0.76, 2.46] | 3.71 | < .001 |
| ot1 x Trial Type | 5.13 | 1.21 | [2.76, 7.50] | 4.25 | < .001 |
| ot2 x Trial Type | -4.17 | 1.21 | [-6.54, -1.80] | -3.45 | < .001 |
| ot3 x Trial Type | -3.65 | 1.21 | [-6.02, -1.28] | -3.02 | 0.003 |
| ot4 x Trial Type | 1.82 | 1.21 | [-0.55, 4.19] | 1.51 | 0.132 |
| Distance Bin | -0.40 | 0.03 | [-0.45, -0.35] | -15.73 | < .001 |
| ot1 x Distance Bin | -2.31 | 0.31 | [-2.93, -1.70] | -7.42 | < .001 |
| ot2 x Distance Bin | -1.22 | 0.31 | [-1.83, -0.61] | -3.90 | < .001 |
| ot3 x Distance Bin | 1.10 | 0.31 | [0.49, 1.71] | 3.53 | < .001 |
| ot4 x Distance Bin | 1.92 | 0.31 | [1.30, 2.53] | 6.14 | < .001 |
| Distance Bin x Trial Type | -0.02 | 0.04 | [-0.09, 0.05] | -0.43 | 0.666 |
| ot1 x Distance Bin x Trial Type | 1.70 | 0.44 | [0.84, 2.56] | 3.85 | < .001 |
| ot2 x Distance Bin x Trial Type | 1.01 | 0.44 | [0.14, 1.87] | 2.28 | 0.022 |
| ot3 x Distance Bin x Trial Type | -1.01 | 0.44 | [-1.87, -0.15] | -2.29 | 0.022 |
| ot4 x Distance Bin x Trial Type | -1.84 | 0.44 | [-2.70, -0.97] | -4.16 | < .001 |

Growth curve formula: $\text{lmer}(\text{Pupil} \sim (\text{ot1} + \text{ot2} + \text{ot3} + \text{ot4}) * \text{Distance Bin} * \text{Trial Type} + (\text{ot1} + \text{ot2} + \text{ot3} + \text{ot4}) / \text{participant}) + (1 / \text{participant} : \text{Trial Type})$ , $\text{control} = \text{lmerControl}(\text{optimizer} = \text{"bobyqa"})$ . Orthogonal polynomial terms: ot1 = linear; ot2 = quadratic; ot3 = cubic; ot4 = quartic.

### Delaying Trial Events Did Not Impact Learning Performance

We examined the extent to which delaying trial events impacted category learning performance and the time participants took to respond when prompted to provide a category decision. A generalized linear mixed effects model of trial-by-trial accuracies (Fig. 2A) revealed that accuracies improved across the learning task (β = .243, *z* = 3.672, *p* < .001). However, the trial-by-trial increase in accuracies across the task did not differ between delayed and immediate trials (β = -.056, *z* = -0.600, *p* = .548). Additionally, overall accuracies on the task were not different between immediate and delayed trials (β = .045, *z* = 0.363, *p* = .716). These results suggest that learning performance was not impacted by delaying trial events compared to the immediate timing that is traditional to learning paradigms. Furthermore, average block accuracies on immediate trials were strongly correlated with accuracies on delayed trials in block 1 (*r* = .55, *p* = .017) and block 2 (*r* = .52, *p* = .025) of learning but were not significantly correlated in block 3 (*r* = .33, *p* = .176) or block 4 (*r* = .29, *p* = .244).

A linear mixed effects model of response times that were time-locked to the ‘Which Category?’ response prompt (Fig. 3B) showed that response times were longer on immediate trials relative to delayed trials (β = .889, *t* = 10.040, *p* < .001) and became faster (i.e., decreased) with task progression (β = -.092, *t* = -5.893, *p* < .001). However, the rate of decrease in response times across the task was not different between delayed and immediate trials (β = .018, *t* = 0.816, *p* = 415). Response times on immediate trials were strongly correlated with response times on delayed trials in all four learning blocks. This indicates that individuals who were fast to respond on immediate trials were also fast to respond on delayed trials. Thus, while response times were longer for immediate trials, delaying trial events with the goal of improving pupil interpretability, did not impact category learning performance.

**Figure 3.**
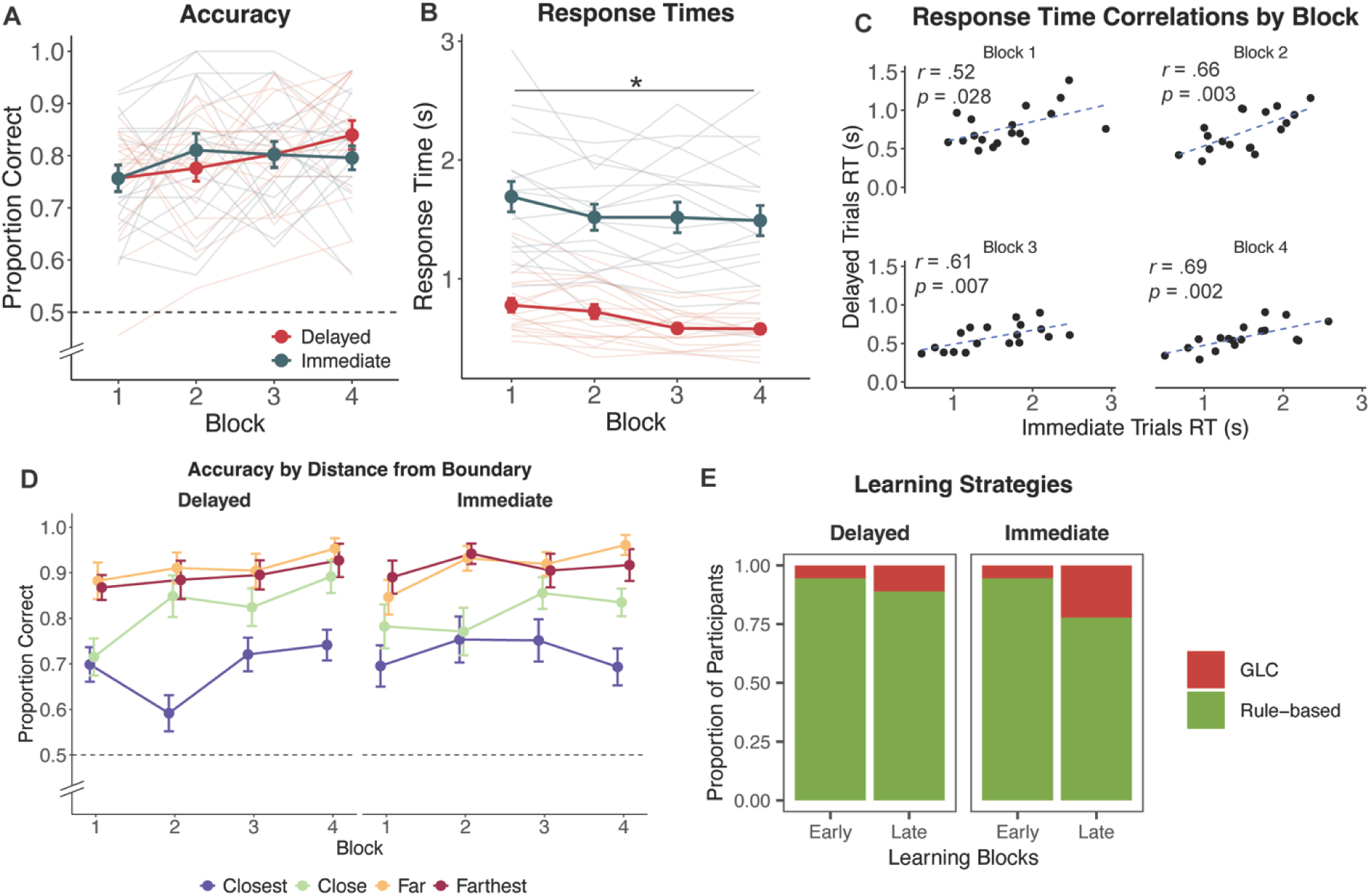
Behavioral outcomes between immediate and delayed trials. A) Accuracies between immediate and delayed trials were not different throughout learning. B) Response times time-locked to the onset of the category response prompt on immediate trials were significantly longer than response times on delayed trials. C) Correlations between response times on delayed trials and immediate trials were significant in all blocks. D) Block accuracies in delayed and immediate trials for stimuli based on distance from category boundary, where stimuli closest are considered to be more difficult to learn than stimuli farthest from the boundary. E) The proportion of participants using the optimal rule-based learning strategy strategy in early blocks (1-2) and in late blocks (3-4) were not different between delayed and immediate trials. A, B, D) Solid lines indicate group average for immediate and delayed trials with standard error of the mean for error bars. Dashed line in A and D denote chance-level.

We also examined the extent to which trial type impacted category learning performance based on stimulus difficulty. Specifically, we were interested in any overall differences in accuracy between trial types and any differences in the rate of learning across the task between stimulus difficulty and trial types. We observed a significant effect of the far (β = .295, *t* = 4.268, *p* < .001) and farthest (β = .296, *t* = 4.287, *p* < .001) stimulus bins on accuracy relative to the closest bin to the optimal category boundary (Table 4). These effects suggest that stimuli further from the category boundary were easier to learn (i.e., higher accuracies) than stimuli closest to the boundary. However, the trial-by-trial increase in accuracy across the task for each stimulus bin was not different between immediate and delayed trials. Therefore, delaying trial events did not impact learning performance based on stimulus difficulty.

**Table 4.** Fixed-effect estimates for linear mixed-effects model of accuracies to examine the effect of trial type, learning block, and stimulus distance from category boundary (observations = 576, groups: participant x block = 72, participant = 18).

| Fixed Effect | Estimate | SE | 95% CI | *t* | *p* |
| --- | --- | --- | --- | --- | --- |
| Intercept | 0.55 | 0.05 | [ 0.44, 0.66] | 10.23 | < .001 |
| Close Bin | 0.12 | 0.07 | [-0.01, 0.26] | 1.76 | 0.078 |
| Far Bin | 0.30 | 0.07 | [ 0.16, 0.43] | 4.27 | < .001 |
| Farthest Bin | 0.30 | 0.07 | [ 0.16, 0.43] | 4.29 | < .001 |
| Block | 0.03 | 0.02 | [ 0.00, 0.07] | 1.93 | 0.054 |
| Trial Type (Delayed vs. Immediate) | 0.13 | 0.07 | [-0.01, 0.27] | 1.88 | 0.060 |
| Block x Close Bin | 0.01 | 0.03 | [-0.04, 0.06] | 0.57 | 0.570 |
| Block x Far Bin | -0.02 | 0.03 | [-0.07, 0.03] | -0.63 | 0.531 |
| Block x Farthest Bin | -0.02 | 0.03 | [-0.07, 0.03] | -0.62 | 0.534 |
| Trial Type x Block | -0.04 | 0.03 | [-0.09, 0.01] | -1.65 | 0.101 |
| Trial Type x Close Bin | -0.13 | 0.10 | [-0.32, 0.07] | -1.30 | 0.196 |
| Trial Type x Far Bin | -0.12 | 0.10 | [-0.32, 0.07] | -1.27 | 0.204 |
| Trial Type x Farthest Bin | -0.07 | 0.10 | [-0.27, 0.12] | -0.76 | 0.446 |
| Trial Type x Block x Close Bin | 0.04 | 0.04 | [-0.03, 0.11] | 1.03 | 0.303 |
| Trial Type x Block x Far Bin | 0.04 | 0.04 | [-0.03, 0.11] | 1.23 | 0.219 |
| Trial Type x Block x Farthest Bin | 0.03 | 0.04 | [-0.04, 0.10] | 0.75 | 0.452 |

Finally, we examined the types of strategies participants used to learn the stimulus categories in early (blocks 1-2) and late (blocks 3-4) learning on delayed trials versus immediate trials with decision-bound computational models (Fig. 2F). In early training, there were no differences in strategy usage between immediate (rule-based: *n* = 17; GLC: *n* = 1) and delayed trials (rule-based: *n* = 17; GLC: *n* = 1; Fisher’s Exact Test, *p* = 1). In late training, there were sixteen rule-based and two GLC strategy users on delayed trials and fourteen rule-based and four GLC strategy users on immediate trials, but the difference between delayed and immediate trials was not significant (Fisher’s Exact Test, *p* = .658). Therefore, participants were using similar strategies to categorize stimuli on immediate trials and delayed trials.

Collectively, the evidence from trial-by-trial accuracies across the task, response times, accuracies based on stimulus difficulty, and learning strategies demonstrate that delaying trial events did not impact learning performance nor the strategies participants used to learn the stimulus categories.

### Decisional Processes

We used DDMs to examine how listeners accumulated sensory evidence (Fig. 4A) and how cautious listeners were to respond (Fig. 4B) on delayed trials across the learning task. The DDMs are Bayesian parameters, therefore, differences between learning blocks can be inferred when the 95% credible intervals are nonoverlapping. Evidence accumulation rates increased by the last block of learning (Mean = 1.771, 95% CI [1.570, 1.985]) relative to the first block (Mean = 1.177, 95% CI [1.015, 1.333]), suggesting that participants became more efficient at extracting information from the stimulus to make a decision as learning progressed. Decision thresholds also changed with learning. Specifically, thresholds were lower in block 4 (Mean = 0.709, 95% CI [0.656, 0.766]) compared to block 1 (Mean = 0.833, 95% CI [0.770, 0.893]), indicating that participants were less cautious to respond as learning progressed.

**Figure 4.**
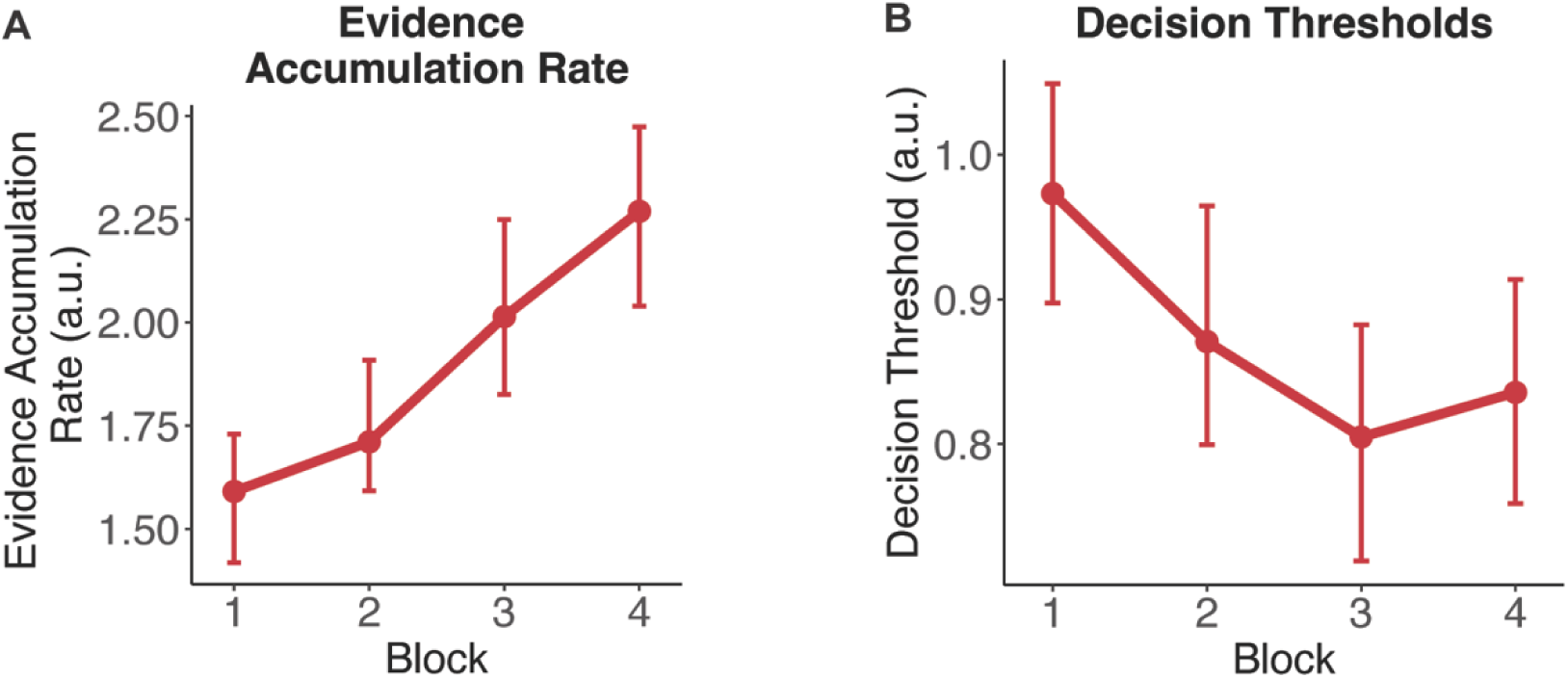
Drift-diffusion model parameters estimated from trial outcomes and response times on delayed trials. Estimated posterior means for A) evidence accumulation rates and B) decision thresholds. Error bars reflect 95% pointwise credible intervals. Differences across blocks can be inferred when the 95% pointwise credible intervals are non-overlapping. Evidence accumulation rates at the end of training (block 4) were higher than the beginning of training (block 1). Decision thresholds at the end of training do not differ from beginning of training.

### Pupil Response Reflects Decisional Processes During Learning

To examine the extent to which the above decisional processes were reflected in the pupillary response, we estimated a GCA on pupillary responses from delayed trials as a function of evidence accumulation rates (Table 5). As evidence accumulation rates increased (i.e., more efficient accumulation), pupillary responses showed significant changes in the intercept, linear, and cubic time terms. Specifically, higher evidence accumulation rates were associated with a smaller pupillary response, a slower rate of dilation, and a stronger secondary curvature.

**Table 5.**
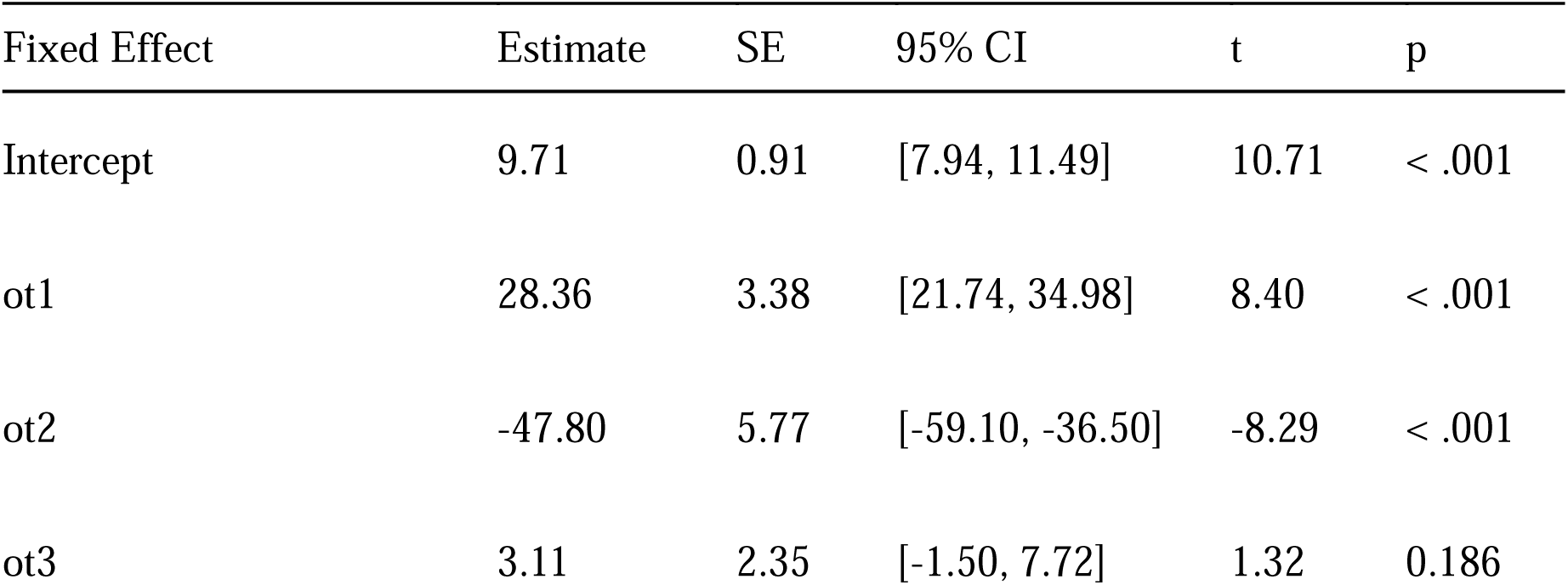
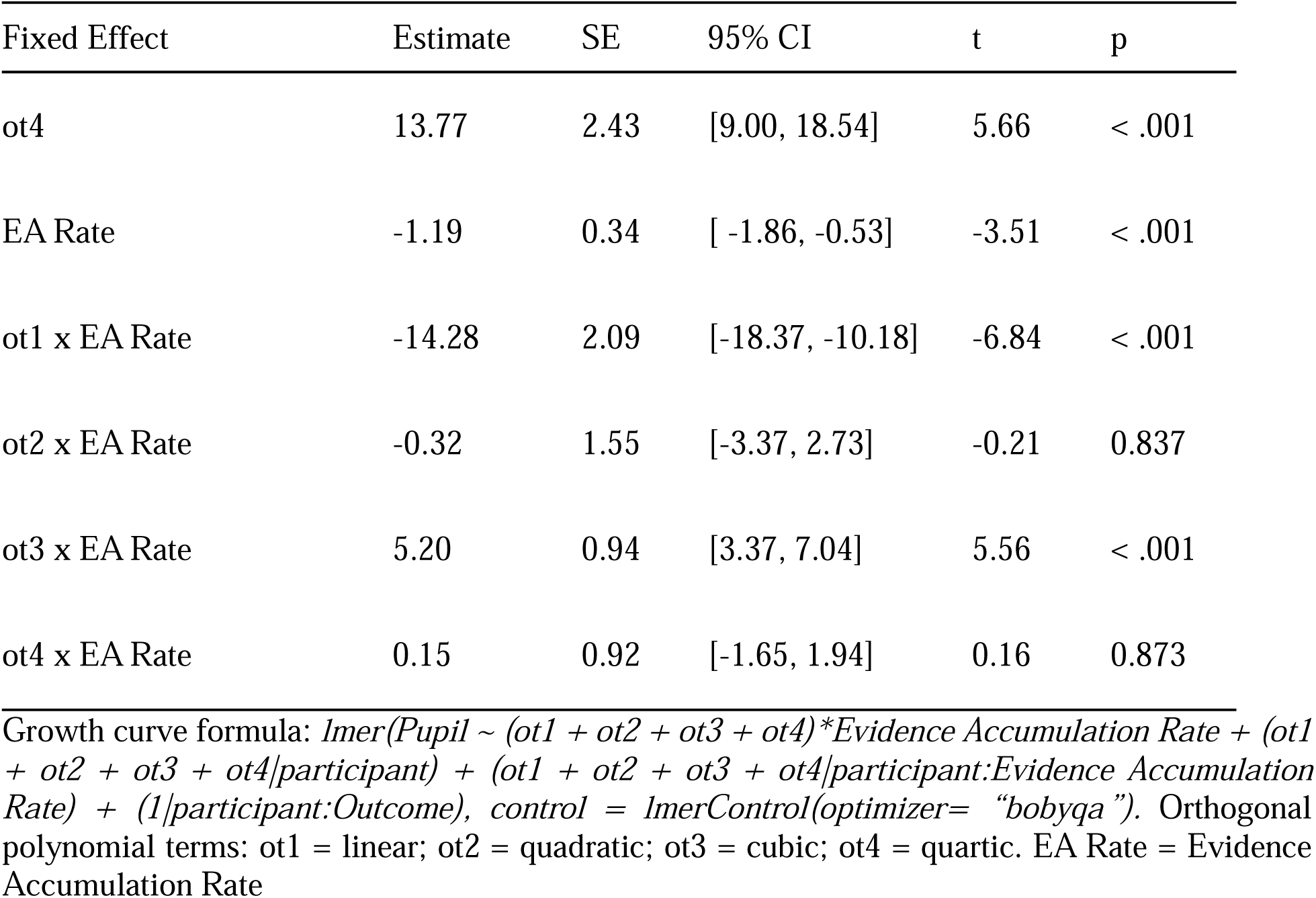
Fixed-effect estimates for model of pupillary responses from 0 to 3000 ms time-locked to stimulus onset to examine the effect of evidence accumulation rates (observations = 5,436, groups: participant x Evidence Accumulation Rate = 36, participant = 18).

We also estimated a similar GCA as a function of decision thresholds (Table 6). Similar to evidence accumulation, there were significant effects of higher decision thresholds (i.e., greater response caution) on the intercept, linear, and cubic time terms. In contrast to evidence accumulation rates, however, greater response caution was associated with a larger pupil response, a faster rate of dilation, and a flatter secondary curvature. Taken together, these findings demonstrate that DDM parameters of decisional processes during learning are reflected in the pupillary response.

**Table 6.**
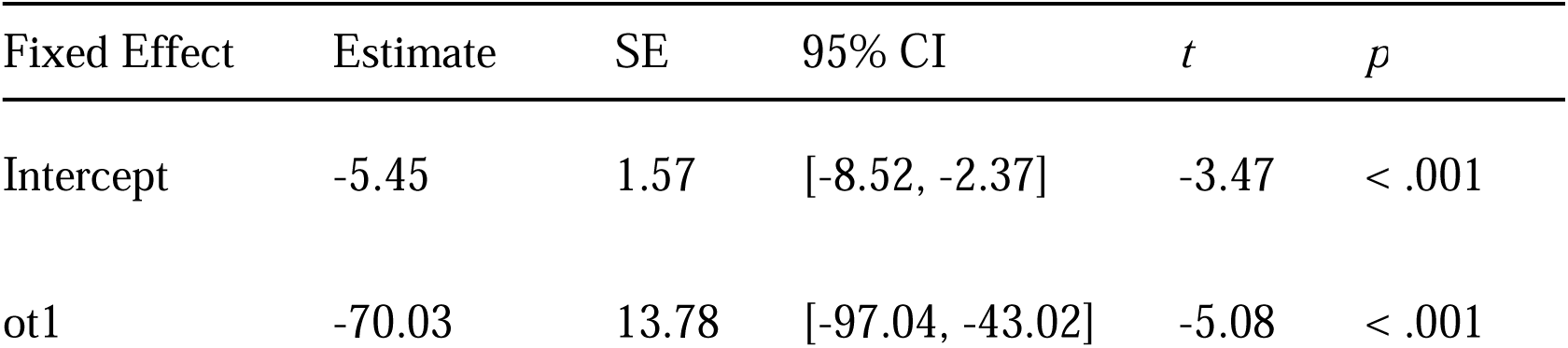
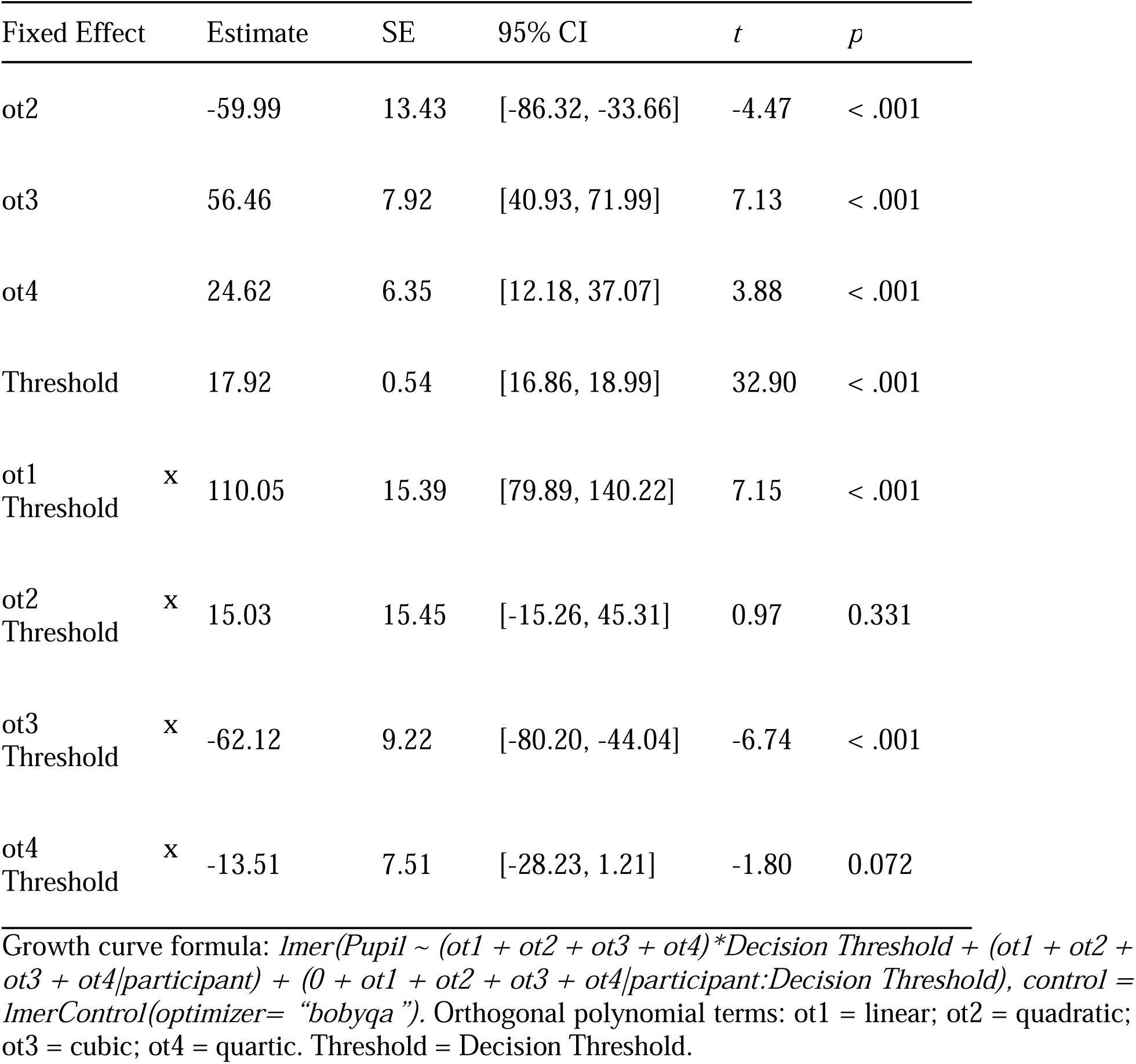
Fixed-effect estimates for model of pupillary responses from 0 to 3000 ms time-locked to stimulus onset to examine the effect of decision thresholds (observations = 5,436, groups: participant x Decision Threshold = 36, participant = 18).

## Discussion

We examined the dynamic processes underlying learning while adult listeners learned to categorize sounds. Importantly, we incorporated pupillometry and DDMs to obtain meaningful insights into the decisional processes involved in learning. Critical to our study design, we implemented timing delays on half of the learning trials to increase the interpretability of the pupil response, while the other half of trials were presented in an immediate manner, which is standard with learning tasks. We found that listeners successfully learned to categorize the sounds, even when trial events were delayed. However, timing delays altered the trajectory of the pupil response, with pupillary responses on delayed trials showing more robust markers of learning, relative to immediate trials. Moreover, decisional processes during learning showed distinct relationships with pupillary responses on these delayed trials. Our results suggest that pupillary responses can be used as a window into the decisional processes that support learning, with minimal information loss when using delayed trials.

Pupillary responses time-locked to stimulus onset dynamically differed in several aspects based on the timing of trial events. When response selection was delayed by four seconds after stimulus presentation, the pupil response showed greater differentiation between correct and incorrect trials, a larger decrease with task progression, and greater differences based on stimulus difficulty, as defined by stimulus distance from the categorical boundary. While some differences in the pupil response were also present when response selection immediately followed stimulus presentation on immediate trials, these differences were less robust than when response selection was delayed. Specifically, the downward slope of the pupil response on immediate trials was relatively stable but showed distinct changes on delayed trials. On average, the pupil response reached its peak prior to the two-second mark and then constricted towards baseline, regardless of trial event timing. On immediate trials, this constriction period comprised the response selection, which required a physical motor execution. The general trend for constriction followed a linear manner on rapid trials. In contrast, the constriction period on delayed trials tended to constrict in a non-linear manner. Moreover, this constriction period provided a ‘clean’ read out of the pupil response that was devoid of motor contaminations, which are known to impact pupillary responses (McGinley et al., 2015; Nelson & Mooney, 2016; Vinck et al., 2015; Yokoi & Weiler, 2022). The absence of motor contamination in the pupil response on delayed trials suggests that the initial dilation period may correspond to sound encoding, while the period after encoding may reflect the underlying cognitive and decisional processing in auditory learning.

The pupillary responses on delayed trials showed distinct markers of learning progress. Consistent with a prior study (McHaney et al., 2021), pupillary responses on delayed trials that were categorized correctly showed a stronger curvature towards baseline than on trials categorized incorrectly. This finding is interesting in that a physical response selection had not yet been initiated, therefore no feedback had been provided. Yet, the pupillary responses suggest that the listener may have already known the outcome of their categorization decision. A similar, stronger curvature in the pupil response was observed with task progression as learning accuracies increased, and the curvature was also stronger for stimuli further from the categorical boundary, which were considered easier to categorize. Taken together, these findings indicate that the downward curvature of the pupillary response after sound encoding may serve as a direct biological marker of the learning process that is optimally captured when the response selection is delayed. We therefore recommend that researchers include timing delays to optimize the pupillary response to better understand learning dynamics.

One aspect of the learning process involves deciding how to map the highly variable auditory stimuli to discrete categorical representations. We examined aspects of this decision-making process during our auditory category learning task using DDMs. DDMs help to describe optimal performance on behavioral tasks and traditionally require trial outcomes and precise response times to derive parameters such as evidence accumulation rates and decision thresholds. One of the major goals of this study was to understand the extent to which accuracies and response times on delayed trials, where response times were conflated by the delay in response selection, could be used to reliably estimate DDM parameters. While accuracies on immediate and delayed trials were not different, response times were significantly faster on delayed trials. Listeners had additional time to processes stimulus information before making a response selection on delayed trials, which may have led to a faster physical response once prompted. Response times were strongly correlated between immediate and delayed trials, which indicated that listeners who were fast to respond on immediate trials were also fast to respond on delayed trials. Our DDM results from trial outcomes and response times on delayed trials indicated that listeners became more efficient at extracting relevant information from the stimulus needed for categorization and required less information to make their decision as they progressed through the task. This finding is consistent with prior research demonstrating an increase in evidence accumulation rates with more training blocks in a speech category learning task (Paulon et al., 2020) and a decrease in decision thresholds during auditory category learning (Roark et al., 2021).

The pupil, behavioral, and computational findings from our study have significant implications for understanding some of the neural mechanisms involved in learning. Specifically, delaying trial events to capture a ‘clean’ pupil response does not impact learning performance in a rule-based auditory category learning task and still allowed for estimation of DDM parameters of decisional processes. It should be noted, however, that our findings may be specific to rule-based category structures. Learning is not impacted by timing delays in trial-by-trial feedback when categories follow a rule-based structure (Chandrasekaran et al., 2014; Maddox et al., 2003). In contrast, categories that are split by non-verbalize rules and require complex integration across acoustic dimensions are known to be impacted by delays in feedback presentation (Chandrasekaran et al., 2014; Maddox et al., 2003). Similar pupillary markers of learning progress and efficacy of DDM parameters estimated from delayed trials may not pan out for category structures that follow a complex integration across multiple acoustic dimensions and requires further investigation.

Changes in pupil size and performance optimization, which can be captured by DDMs, share common mechanisms involving the locus coeruleus-norepinephrine system (Gold & Shadlen, 2000, 2002, 2007; Mulder et al., 2014; Shadlen & Newsome, 2001) and basal forebrain cholinergic system (Bentley et al., 2011; Mridha et al., 2021; Reimer et al., 2016). Therefore, pupillary responses during learning should reflect parameters of decision-making derived from the DDM. Prior studies have demonstrated relationships between baseline pupil diameter and metrics of decision-making (de Gee et al., 2014, 2020, 2023; Nassar et al., 2012), and difficult decisions being associated with larger pupil dilation and higher decision thresholds (Cavanagh et al., 2014). Here, we examined task-evoked pupillary responses during learning and observed significant changes in the pupil response based on learners’ evidence accumulation rates and decision thresholds. Specifically, the secondary curvature of the pupil response was sharper for learners who were more efficient at extracting information from the stimulus and for those who were less cautious responders. Our findings indicate that task-evoked pupillary responses may serve as a window into online decisional processes during learning.

With the identification of physiological markers of learning success in the pupillary response, researchers can begin targeting the mechanisms underlying learning and leverage these biological markers as metrics of intervention efficacy. For instance, vagus nerve stimulation is growing in popularity for improving task performance and has been shown to enhance learning of novel speech categories (Llanos et al., 2020; McHaney et al., 2023). Behavioral enhancement induced by vagus nerve stimulation is hypothesized to operate via the LC-NE and cholinergic systems (Hulsey et al., 2016; Schuerman et al., 2022; Vonck & Larsen, 2018), which again, is a shared mechanism for performance optimization measured by DDMs and changes in pupil size. The reason for improved learning of novel speech categories with vagus nerve stimulation in prior studies remains unclear, but future studies could leverage the integration of DDMs and pupillometry to assess the extent to which learners become more efficient evidence accumulators and/or less cautious responders with the application of vagus nerve stimulation. These findings would help to inform why some learners do better than others, from a mechanistic standpoint.

Our study sought not only to elucidate the underlying mechanisms of auditory category learning but also to provide recommendations for a methodological approach that could be applied to a broader spectrum of cognitive-behavioral tasks. Crucially, our study demonstrated that delaying trial events to increase the interpretability of the pupil response that was free of motor confounds did not impact learning performance. Response times between immediate and delayed trials were strongly correlated, hence DDM parameters estimated from delayed trials were likely meaningful and interpretable. By delaying trial events to optimize pupillary responses, we found that the curvature of the pupil response after sound encoding, but prior to response selection, was indicative of cognitive processing that related to trial outcome, stimulus difficulty, and crucially, decision-making processes. We recommend timing delays should be incorporated to increase the interpretability of cognitive and decisional processes from pupillary responses. It is important to acknowledge that immediate feedback timing is critical to learning performance when stimulus category structures require complex integration across dimensions. Future studies should examine the extent to which these pupillary markers of learning and cognitive-decisional processing hold true for such stimuli with more complex category structures to fully understand the learning and perceptual dynamics that can be interpreted from pupillary responses.

## Data and Code Availability

The code and datasets generated and/or analyzed during the current study are available in the Open Science Framework repository and can be accessed at https://osf.io/a9grj/.

## CRediT Authorship

Jacie R. McHaney: Conceptualization, Methodology, Software, Formal Analysis, Data Curation, Writing – Original Draft, Visualization, Supervision, Project Administration; Casey L. Roark: Conceptualization, Methodology, Resources, Writing – Review & Editing; Matthew J. McGinley: Conceptualization, Methodology, Writing – Review & Editing, Funding Acquisition; Bharath Chandrasekaran: Conceptualization, Methodology, Resources, Writing – Original Draft, Supervision, Funding Acquisition

## Author Note

The authors have no conflicts of interest to disclose.

